# Isolation And Identification Of Major Pathogenic Bacteria From Clinical Mastitic Cows In Asella Town, Ethiopia

**DOI:** 10.1101/2020.05.26.118141

**Authors:** Gezehagn Kasa Tufa, Betelhem Tegegne Muluneh, Belege Tadesse Siyamregn

## Abstract

Mastitis is a multi-etiological and complex disease causing inflammation of parenchyma of mammary glands is a problem in many dairy herds. The objective of this study was isolation and identification of the pathogenic bacteria that cause bovine clinical mastitis. A cross sectional study was undertaken from November 2018 to April 2019 on small scale and government dairy farms in Asella town. Cows were examined directly at quarter and teat level for clinical manifestation. A total of 83 milk samples were collected from 46 cows that shows clinical sign of mastitis from a total of 12 farms. Isolation and identification of major bacterial species was carried out by culturing on different media and using primary and secondary biochemical tests. Out of the 83 samples collected and examined, all (100%) were positive for cultural isolation of bacterial species. The bacteria were identified to genus and species level. Among the 83 isolates 32 (38.6%) were *S. aureus*, 24 (28.9%) were *Staphylococcus intermedius* and 6 (7.2%) were *Staphylococcus hyicus*, other bacteria like *Escherichia coli* 12(14.5%), Streptococcus species 2 (2.4%) were also isolated. Bacillus Species 2 (2.4%), Proteus species 2(2.4%) and 3 (3.6%) of them were mixed bacterial infections. The present study revealed that both contagious and environmental bacterial pathogens were responsible for the occurrence of clinical mastitis. Proper milking practices and farm husbandry practices as well as future detailed studies up to the species level and on antibiotic profiles of the pathogens are needed.

## 1. INTRODUCTION

Ethiopia is believed to have the largest livestock population in Africa. The total cattle population in the country is estimated to be about 56.71 million. Out of this, the female cattle constitute about 55.45% and the remaining 44.55% are male cattle. From the total cattle 98.66% in the country are local breeds and the remaining are hybrid and pure exotic breeds that accounted for about 1.19 and 0.14%, respectively (CSA, 2015). The livestock sector has been contributing considerable portion to the economy of the country and still promising to rally round the economic development of the country. However, milk production does not satisfy the country’s requirements due to a multitude of factors (Biffa *et al*., 2005). Mastitis is among the various factors contributing to reduced milk production. Bovine mastitis is the second most frequent disease next to reproductive disorders and one of the major causes for economy failure in Ethiopia. It affects both the quantity and quality of milk (Capuco *et al*., 1992).

Mastitis is a multi-etiological and complex disease, which is defined as inflammation of parenchyma of mammary glands (Radostis *et al*., 2000). The disease mainly resulted from injurius agents including pathogenic microorganisms, trauma and chemical irritants. Even if it occurs due to the injury of any type, the udder disease of major concern is that associated with microbial infection. Among various infectious agents, bacterial pathogens have been known to be widely distributed in the environment of dairy cows, constituting a threat to the mammary gland (Radostits *et al*., 2007). Over 130 different microorganisms have been isolated from mastitis positive cow milk samples, of which almost all are bacteria. The most common pathogens comprise contagious bacteria mainly *Staphylococcus aureus* and *Streptococcus agalactia* and Environmental bacteria mainly coliforms and some species of streptococci that are commonly present in environment (Radostits *et al*., 2007; Quinn *et al*., 2002).

Mastitis can be manifested by a wide range of clinical and subclinical conditions. Clinical mastitis is characterized by sudden onset, alterations of milk composition and appearance, decreased milk production, and the presence of the cardinal signs of inflammation in infected mammary quarters. It is readily superficial and visually detected. It occurs when the inflammatory response is strong enough to cause visible changes in the milk (clots, flakes), the udder (swelling) or the cow (off feed or fever). Even if there is a great loss related with both conditions, clinical mastitis continues to be a problem in many dairy herds (Hogan *et al*., 1999; Harmon, 1994).

Mastitis is a global problem as it adversely affects animal health, quality of milk and the economis of a country by causing huge financial losses (sharm *et al*., 2007). There is agreement among authors that mastitis is the most widespread infectious disease in dairy cattle and from an economic aspect, the most damaging (Tiwari *et al*., 2010). This disease has also been known to cause a great deal of loss or reduction of productivity, to influence the quality and quantity of milk yield and to cause culling of animals at an unacceptable age. Most estimates have shown a 30% reduction in productivity per affected quarter and a 15% reduction in production per cow/lactation, making the disease one of the most costly and serious problems affecting the dairy industry worldwide (Hogan *et al*., 1999). Clinical mastitis in a dairy herd is threatening to a farmer but treatment can be given immediately to control it (Harmon, 1994).Mastitis is worth to study as it incurs financial losses attributed to reduced milk yield, discarded milk following antibiotic therapy, early culling of cows, veterinary costs, drug costs, increased labor, death in per acute septicemia and replacement cost (Nesru *et al*., 1997). In Ethiopia, even though the disease of mastitis has been known locally, it has not been studied systematically (Sori *et al*., 2005). More over clinical mastitis is frequently occurring and economically important disease for dairy industry in our country, Ethiopia. Regardless of this, very little attention is also given to mastitis in the country and efforts have only been concentrated on the treatment of clinical cases. In addition to this for the control and prevention of the disease, proper isolation and identification of the responsible bacterial agents is necessary regarding to which little studies are still done. Therefore this study was done to isolate and identify pathogenic bacteria from cows that have clinical mastitis.

## 2. MATERIALS AND METHODS

### 2.1. Study Area

The study was conducted from November 2018 to April 2019 in Asella town. The town is located in Arsi Zone of Oromia region about 175 km from Addis Ababa. The town has a latitude and longitude of 7°57′N 39°7′E, with an elevation of 2,430 meters above sea level. Topographically Asella is a highland area with annual rain fall of 2300 to 2400mm (ATAO, 2016).

### 2.2. Study Animals

Lactating dairy cows found in privately owned small holder dairy farms and government dairy farms in Asella town were involved in the study population. The study was conducted on purposely selected lactating dairy cows with clinical sign of illness regardless of the age, breed, pregnancy, husbandry system, hygienic condition, milking practice, parity and stage of lactation.

### 2.3. Study Design and sampling

A cross sectional study using laboratory isolation and identification of bacteria was undertaken from November 2018 to April 2019 on small scale and government owned dairy farm in Asella town. Cows that shows signs of clinical mastitis were selected and sampled purposively from the farms that are found in the town (all dairy farms found in the town were included in the study).

### 2.4. Data Collection

#### 2.4.1. Questionnaire survey

The data about the factors like age, breed, pregnancy, parity number and lactation stages of the cows and milking practice, husbandry system and hygienic condition of the farms were collected from owners and farm managers through a face to face questionnaire survey.

#### 2.4.2. Physical examination of udder and milk

Cows were examined at quarter and teat level for the observation of signs of clinical mastitis. The udders of the study cows were examined visually and by palpation for the presence of clinical mastitis. During examination attention was paid to cardinal signs of inflammation, size and consistency of udder quarters. Inspection of milk for discoloration, consistency and presence of clots, which are characteristics of clinical mastitis were performed.

#### 2.4.3. Milk sample collection

A total of 83 milk samples were collected from 46 cows which show clinical signs of mastitis in Asella town from a total of 12 small scales and government owned dairy farms. The milk samples were collected from the teats of clinically infected quarter’s from cows that are not treated early with either intra mammary or systematic antimicrobial agents. Milk sampling was carried out following aseptic procedures as described by National Mastitis Council (2004).

##### Collection, transportation and storage of milk samples

Udder was first washed with water and then the teats and teat orifices were disinfected with pieces of cotton wool soaked in 70% ethyl alcohol and then dried with fresh pieces of cotton wool. Approximately 5-6 ml of milk from infected quarter were taken (after discarding the fore milk) aseptically in sterile bottles for bacteriological investigation and labeled. Samples were placed in ice box containing ice packs and transport immediately to microbiology room of Asella Regional Veterinary laboratory. Samples which are not processed immediately were preserved in refrigerator at 4°C until processing.

### 2.5. Laboratory Diagnosis

#### 2.5.1. Bacterial isolation

For the isolation of bacteria, different types of media were used (solid and liquid media). The common media used during the study were blood agar, nutrient agar, MacConkey agar, manitol salt agar, Eosin methylene blue medium, nutrient broth, Triple sugar iron agar, Simon citrate agar, Tryptophan broth and MR-VP broth biochemical media were used.

##### Preparation of culture media

To prepare the media for bacterial culture, the manufacturer’s instructions were followed. All glasses, flasks and petridishes used for the preparation of media were first sterilized using appropriate sterilizers like autoclave. The appropriate amounts of dehydrated media were weighed using sensitive balance and the required amount of distilled water was added to the agar media powder in the flask. Then they were dissolved in heating mantle until it boiled and frothy appearance was settled (removed), then the media were sterilized by autoclave at 121^0^C for 15 min holding time, and cooled in water bath at 50^0^C before dispensed in to the petridishes. Some media like blood agar requires addition of blood after it is cooled to 50^0^C since RBC do not tolerate higher temperature (Quinn *et al*., 2002).

##### Cultural methods

The samples collected from cows were cultured on general purpose media such as blood agar and nutrient agar using sterile loop inside the biosafety cabinet and around Bunsen burner. Other selective and differential media such as Manitol salt agar, MacConkey agar, Eosin metheylne blue agar were also used for cultural purpose. The media were incubated aerobically at 37ºC for 24 hours.

##### Examination of culture

Visual examination was done for detection of growth, pigmentation, haemolysis and colonial morphology.

#### 2.5.2. Identification of isolates

##### Gram’s staining

It was used to study morphology, shape and gram staining reaction of each isolates. Gram-positive bacteria appeared purple, while Gram-negative bacteria appeared red.

##### Biochemical tests

The staining is followed by use of various biochemical reagents and tests to get closer to the identification of bacteria. There are many biochemical tests available for bacterial identification.

Primary biochemical tests: used to identify organisms into genus level. The primary biochemical tests that were applied are Catalase test andOxidase tests.

Secondary biochemical tests: used to identify organism into species level. Secondary biochemical tests performed includes Indole test, Methyl (MR) test, Voges–Proskauer (VP) test, Citrate utilization test, Triple sugar iron test and coagulase test.

### 2.6. Data Management and Analysis

The data including the quarter affected, parity no, husbandry system, hygienic condition, lactation stage and milking practice were recorded depending on clinical inspection; pathogenic bacteria isolated and identified were entered into Microsoft Excel computer program 2007. STATA version 14was used to summarize the data and descriptive statistics like percentages were used to express the result.

## 3. RESULTS

### 3.1. Bacteria Isolated from Mastitic Milk

In the current study all (100%) the 83 milk samples collected from clinically mastitic cow were positive for cultural isolation of bacterial species. The bacteria were also identified to genus and species level. Most isolates were Staphylococcus species 62 (74.7%) in which all of them were coagulase positive. Among the total of 83 isolates 32 (38.6%) were *Staphylococcus aureus*, 24 (28.9%) were *Staphylococcus intermedius* and 6 (7.2%) were *Staphylococcus hyicus*. Other bacteria like *E.coli* 12 (14.5%), Streptococcus species 2 (2.4%) were also isolated (Table 1).

**Table 1:** Frequency and percentage of various bacterial species isolated from clinical mastitic samples

| Bacterial species | Frequency | Percentage (%) |
| --- | --- | --- |
| *S.aureus* | 32 | 38.6 |
| *S.intermedius* | 24 | 28.9 |
| *S.hyicus* | 6 | 7.2 |
| *E.coli* | 12 | 14.5 |
| Bacillus species | 2 | 2.4 |
| Protes species | 2 | 2.4 |
| Streptococcus species | 2 | 2.4 |
| Mixed infection | 3 | 3.6 |
| Total | 83 | 100 |

#### 3.1.1. Clinical mastitis at teat level in different quarters

There was no significant difference between quarters in the occurrence of pathogenic microorganisms (p>0.05). However, the highest proportion of microorganisms has been isolated from left back teat, 25(30.1%). From all teats except left front, the highest proportion has been recorded in Staphylococcus aureus (Table 2).

**Table 21:**
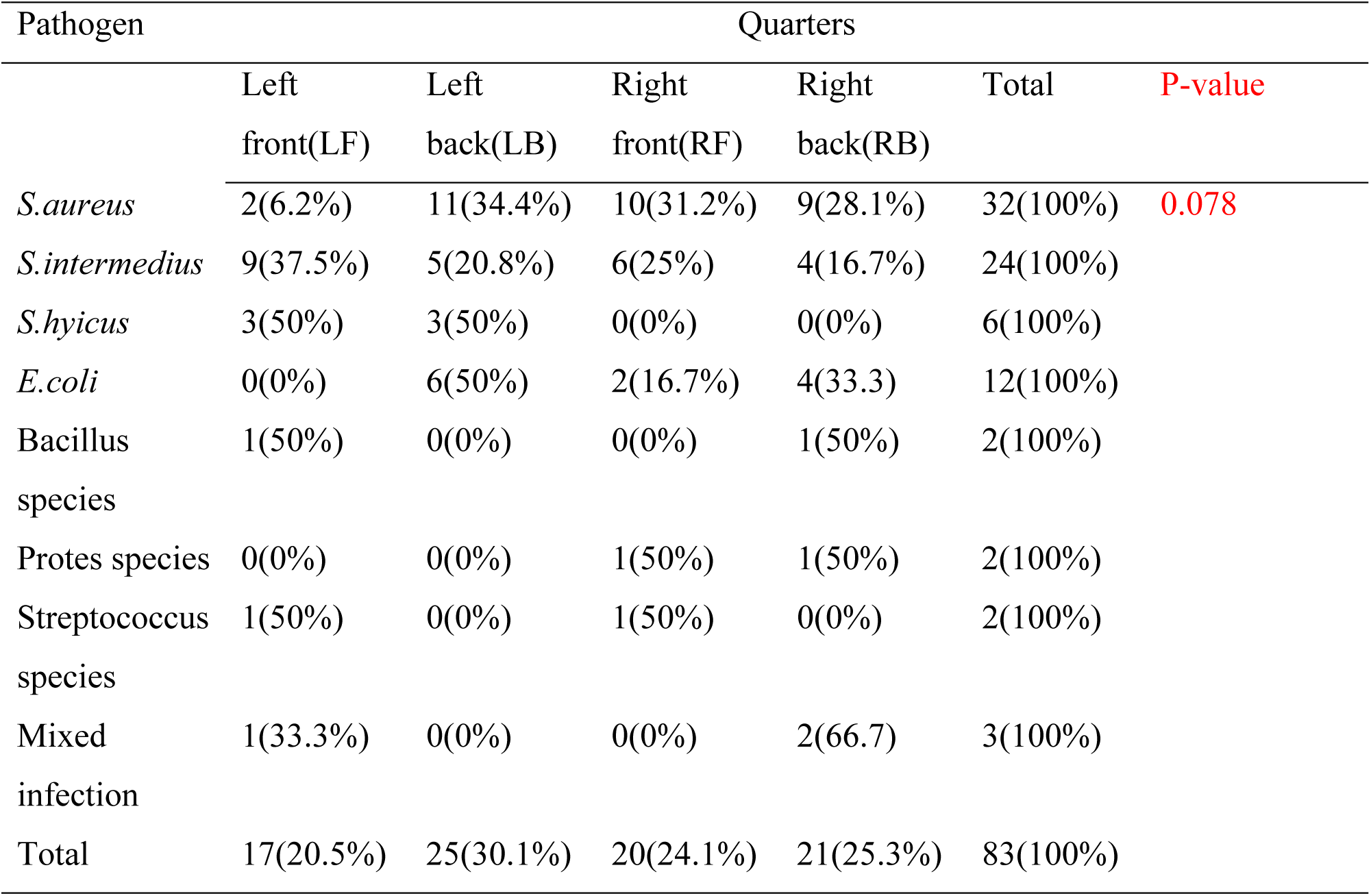
Proportion/infection rate of clinical mastitis at teat level in different quarters

**Table 2:** Biochemical reactions of gram positive isolated organisms.

| Bacterial<br>species | Biochemical tests |  |  |  |  |  |
| --- | --- | --- | --- | --- | --- | --- |
|  | Gram<br>reaction | Color | Shape | Catalase | Oxidase | Coagulase |
| *S. aureus* | + | Purple | Cocci<br>(bunches of<br>grape) | + | - | + |
| *S.intermedius* | + | Purple | Cocci<br>(bunches of<br>grape) | + | - | + |
| *S.hyicus* | + | Purple | Cocci<br>(bunches of<br>grape) | + | - | + |
| Streptococcus<br>species | + | Purple | Chain<br>of Cocci | - | - | - |
| Bacillus<br>species | + | Purple | Long road | + | + | - |

**Table 3:**
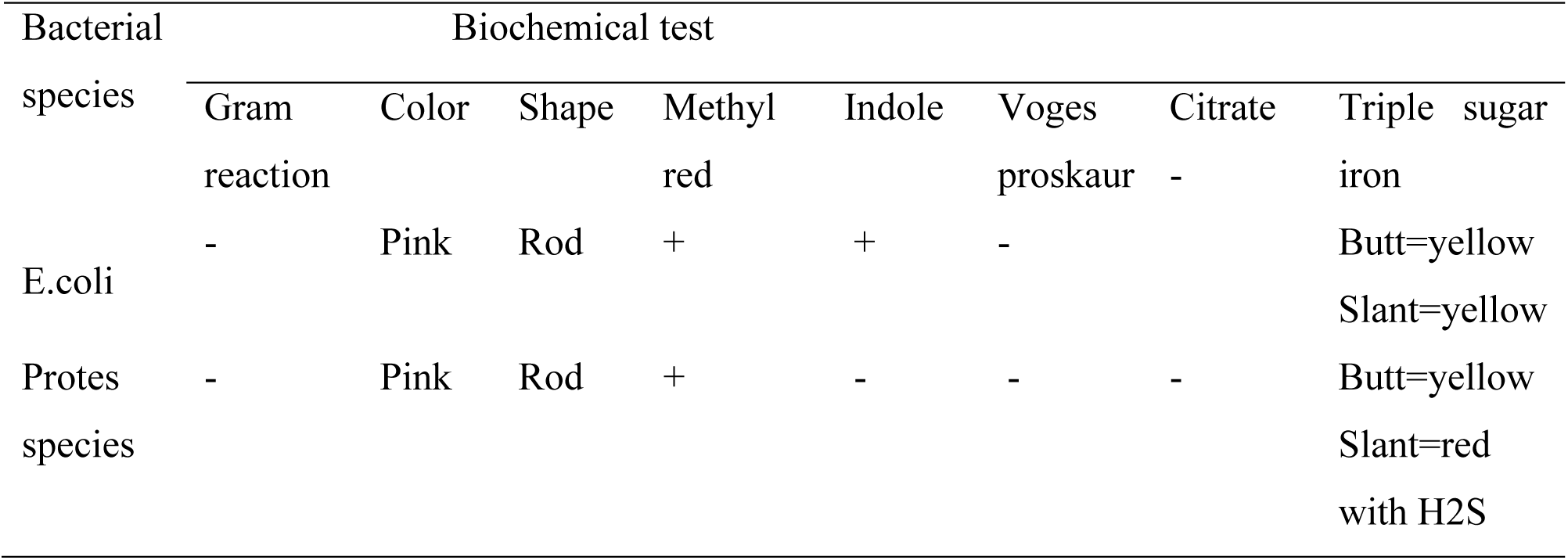
Biochemical reactions of gram negative isolated organisms.

| Bacterial species | Biochemical test |  |  |  |  |  |  |  |
| --- | --- | --- | --- | --- | --- | --- | --- | --- |
|  | Gram reaction | Color | Shape | Methyl red | Indole | Voges proskaur | Citrate - | Triple sugar iron |
| E.coli | - | Pink | Rod | + | + | - |  | Butt=yellow<br>Slant=yellow |
| Protes species | - | Pink | Rod | + | - | - | - | Butt=yellow<br>Slant=red<br>with H2S |

### 4.2. Results of the Questionnaire Survey

The study was performed in 12 farms and out of them 10(83.3%) farms were managed intensively and the other 2(16.7%) were semi-intensive farms. From the total number of farms in which the study was conducted 6(50%) of the farms were managed in a poor hygienic condition. From the farms included in the study 11(91.2%) farms kept only cross breed cows, whereas only one farm kept local breed cows (Table 3).

**Table 3:** Farm level description of factors

| Variable |  | Frequency | Percentage |
| --- | --- | --- | --- |
| Husbandry system | Intensive | 10 | 83.3 |
|  | Semi-intensive | 2 | 16.7 |
| Hygienic condition | Poor | 6 | 50 |
|  | Medium | 4 | 33.3 |
|  | Good | 2 | 16.7 |
| Breed | Local (Borena) | 1 | 8.3 |
|  | Cross | 11 | 91.7 |
| Total farms |  | 12 | 100 |

The proportions of clinical mastitis was high in cows that have high parity numbers, in early lactation stage and in cross breed cows as compared to low parity, late and late lactation stage and local breed cows respectively (Table 4).

**Table 4:** Profiles of cows included in the study

| Variable |  | Frequency | Proportion |
| --- | --- | --- | --- |
| Breed | Local(Borena) | 2 | 4.3 |
|  | Cross | 44 | 95.7 |
| Age | Young | 1 | 2.2 |
|  | Adult | 39 | 84.8 |
|  | Old | 6 | 13 |
| Parity no | 1 | 6 | 13.04 |
|  | 2 | 9 | 19.57 |
|  | 3 | 12 | 26.09 |
|  | 4 | 12 | 26.09 |
|  | 5 | 6 | 13.04 |
|  | 6 | 1 | 2.18 |
| Pregnancy | Pregnant | 17 | 37 |
|  | Non-pregnant | 29 | 63 |
| Lactation stage | Early | 19 | 41.3 |
|  | Mid | 15 | 32.6 |
|  | Late | 12 | 26.1 |
| Total |  | 46 | 100 |

## 4. DISCUSSION

A total of 83 Milk samples were collected and processed from clinically infected cows from small scale holder and government dairy cows in Asella town. The result of the current study showed that staphylococcus species, streptococcus species, *Escherichia coli*, bacillus species and protes species were isolated which has been also reported in other study (Biruke and Shimeles, 2012).

Isolation and identification of pathogenic bacteria such as 38.6% *S. aureus*, 28.9% *S. intermedius*, shows the high contributions of microbial agents as a cause of mastitis in the area. From coagulase positive staphylococcus species the predominant pathogen isolated in current study was *S. aureus* (38.6%). This finding is in agreement with 39.1% reported by Bedada and Hiko (2011) whereas, higher than the report of Mulugeta and Wassie (2013) who reported isolation rate of 30.0%. The reason for higher isolation rates of *S. aureus* is its wide ecological distribution inside the mammary gland and skin. In areas where hand milking and improper use of drug is practiced to treat mastitis case, its dominance has been suggested and might be due to the fact that they are easily transmitted during milking via the milker’s hands as it is contagious pathogens (Jones *et al*., 1998).

Laboratory results from the current study indicated that the prevalence of *Staphylococcus intermedius* was 28.9% which is the second predominant isolated bacteria next to *Staphylococcus aureus*. The result reported from the current study was lower than the (38.4%) reported by Argaw and Tolosa (2008) but much higher than reports of Birhanu *et al*. (2013) who reported 7.14% in the same study site. The variability in the prevalence of isolated bacteria between reports could be attributed to differences in management of the farms, milking practices and hyigienic condition of the farms. (Mungube *et al*., 2005).

The 14.5% isolation rate of *E. coli* found in this study was comparable with the findings of Biruke and Shimeles (2012) who reported 18.6% at Addis Ababa, while it was found to be lower than the 40.7% reported by Iqbal *et al*. (2004 and much higher than the report of Birhanu *et al*. (2013), who reported 5.71% in different parts of Ethiopia. The prevalence of environmental *E. coli* may be associated with poor farm cleanliness and poor slope of stable areas. Faeces which are common sources of *E. coli* can contaminate the premises directly or indirectly through bedding, calving stalls, udder wash water and milker’s hands (Radostits *et al*., 2007).

The 2.4% proportion of streptococcus species found in this study was much lower than the findings Biruke and Shimelis, (2012) who reported 16.7% streptococcus species. The variability in the prevalence of isolated streptococcus species between reports could be because of some contagious Streptococcus species survives poorly outside the udder, and established infections are eliminated by frequent use of penicillin and other antibiotics and because of difference in the milking practice between the different farms in the studies (Radostits *et al*., 2007).

The results fromthe current study indicated that the proportions of bacillus species were low (2.4%) which was similar with findings of Bedada and Hiko (2011) who reported 3.4% proportion. Bacillus species are only occasionally mastitis causing pathogens. The infection is associated with contamination of teat injures and surgery. The level of infection can be high during the dry period following the use of dry cow therapy preparation which may have been contaminated with the organisms (Radostits *et al*., 2007).

The 2.4% Proteus species isolated was almost similar with 2.63% report of Hussein (1999) in and around Addis Ababa and 2.2% report of Bedada and Hiko (2011). The prevalence of Proteus species might be due to the residing of this agent in the cow’s environment bedding, feed and water. They spread due to poor environmental sanitation and milking practice.

The current study revealed that clinical mastitis has affected cows at different stages of lactation, early (41.3%), mid (32.6%) and late (26.1), which was comparable with the finding of Kerro and Tareke, (2003) who reported a high prevalence rate of clinical mastitis of cow in early lactation.

The occurrence of mastitis for cows that gives birth for 3^rd^ and 4^th^ times was 26.09%; which was lower than the findings of Biruke and Shimelis (2012) who reported a prevalence of 71.5% during 3^rd^ and 4^th^ parity. In this study during mid parity number a high proportion was recorded. This could be associated with the possibility of exposure to the infectious agent with increasing number of parity. This was in agreement with the findings of Biffa *et al*. (2005) and Tesfaye (1995). Again it was also agrees with the report of Biruke and Shimelis (2012) who report high proportion during the medium of parity and when reach 5^th^ parity number it reduces, this is due to the farm management system, means culling of too old lactating cow and there is small number of cows giving birth for fifth (5^th^) times and more.

The current study revealed that highest proportion (84.8%) of clinical mastitis was found in adult lactating cows of ages between 3and 9 years, followed by old cows of ages greater than 9 years (13%) and the lowest prevalence (2.2%) was recorded in young cows with ages of 2 years. Increase in occurrence of mastitis with the ages could be due to an increased period of exposure of the udder during previous ages of lactating cows, but because of less number of older cows in the studied farms because of culling of aged lactating cows, the proportions of clinical mastitis in older lactating cows was lower than adult cows. Correspondingly, Teshome *et al*. (2018) reported highest prevalence (48.78%) in lactating cows of ages greater than 8 years, followed by cows of ages 4-8 years (30.54%) and the lowest prevalence (18.52%) was reported in cows of ages less than 4 years. The increase in proportion of mastitis with age might be due to the physiology of exhausted canal which is more dilated and remains partially open due to years of repeated milking. This could have facilitated the entrance of environmental and skin associated microorganisms leading clinical mastitis (Blowey and Edmondson, 2010).

## 5. CONCLUSION AND RECOMMENDATIONS

The present study which was conducted on isolation and identification of major bacterial pathogens from clinically ill cows revealed that both contagious and environmental pathogens, such as *S. aureus, S. intermedius, S. hyicus*, streptococcus species, Bacillus species, protes species, *E. coli* were isolated. From the isolated organisms, *S. aureus (*38.6%), *S. intermedius (*28.9%) and E.coli (14.5%) were the predominant organisms. This indicates that contagious mastitis is one of the major problems of dairy cows in milk production followed by environmental mastitisTo reduce the problem of clinical mastitis proper milking practices like milking of infected cows after milking of apparently healthy animals and regular cleaning of cow’s udder should be practiced. Farm husbandry practices should be maintained to avoid contamination of cows’ house and bedding to control and prevent environmental mastitis. There is a need for further detailed studies on different pathogenic micro organisms to the species level and antibiotic susceptibility pattern of those microorganisms that cause clinical mastitis.

## 7. ANNEXES

**Annex 1:** Questionnaires to assess farm conditions in As sela town

Name of the farm___________

Owner’s name___________ Address___________

Husbandry system

Extensive___________ Semi intensive___________ Intensive___________

I. Housing

1. Type of housing for your milking cows:

Tie-stall___________ Free-stall___________

2. What type of material is the base of your milking cow’s house made off?

Concrete___________ Sand___________ mattress other (specify):___________

3. How are the house cleaned?

Not applicable___________ Scraped___________ flushed with water___________ other (specify):___________

4. How many times per day house is cleaned?

Not applicable___________, ___________times/

5. Do you disinfect the house?

Yes___________ No___________

5.1. If say yes for question 5 how many times disinfect within a month?

___________times/month, Other (specify)___________

6. How often do you clean out manure from your milking cows’ house??

Never___________, ___________times/day, other (specify):___________

II. Milking practice

8. What types of milking practice you use?

Milking machine___________ Hand milking___________

9. Are udders washed or sprayed with water before milking?

Yes___________ No___________

10. Are teats disinfected before milking (pre-dip)?

If you say yes, proceed to 11 question. If No, skip 11 question

11. How is the pre-dip applied?

Sprayed___________ dipped___________

12. Are teats dried before milking?

Yes___________ No___________

13. How do you dry teats prior to milking time?

Not applicable___________ Disposable paper towel (or newspaper)___________

Reusable cloth towel___________ I do not dry teats___________ Other (specify)___________

14. Do you use separate drying material for each cow?

Yes___________ No___________

15. If you use reusable towel, do you wash or disinfect these towels after every milking?

Not applicable___________ Yes___________ No___________

16. Are teats disinfected after milking (post-dip)?

If you say Yes, proceed to 17 question and if No, skip 17 question

17. How is the post-dip applied?

Sprayed___________ Dipped___________

